# Loss of CD24 promotes radiation- and chemo-resistance by inducing stemness properties associated with a hybrid E/M state in breast cancer cells

**DOI:** 10.1101/2022.05.05.490732

**Authors:** Isaline Bontemps, Céline Lallemand, Denis Biard, Nathalie Dechamps, Thierry Kortulewski, Emmanuelle Bourneuf, Capucine Siberchicot, François Boussin, Sylvie Chevillard, Anna Campalans, Jérôme Lebeau

**Affiliations:** Université de Paris and Université Paris-Saclay, LCE/iRCM/IBFJ CEA, Stabilité Génétique Cellules Souches et Radiations, F-92265, Fontenay-aux-Roses, France; Université Paris-Saclay, SEPIA/IBFJ CEA, Team Cellular Engineering and Human Syndromes, F-92265, Fontenay-aux-Roses, France; Université de Paris and Université Paris-Saclay, Inserm, Cytometry platform/iRCM/IBFJ CEA, Stabilité Génétique Cellules Souches et Radiations, F-92265, Fontenay-aux-Roses, France; Université de Paris and Université Paris-Saclay, Inserm, LRP/iRCM/IBFJ CEA, UMR Stabilité Génétique Cellules Souches et Radiations, F-92265, Fontenay-aux-Roses, France

**Keywords:** Breast cancer, CD24, cancer stem cells, EMT, resistance

## Abstract

There is compelling evidence that cancer stem cells (CSCs) play an essential role in failure of conventional antitumor therapy. In breast cancer, CD24^-/low^/CD44^+^ phenotype as well as a high aldehyde dehydrogenase activity (ALDH^+^) are widely associated with CSC subtypes. Furthermore, CD24^-/low^/CD44^+^ pattern is also characteristic of the mesenchymal cells generated by an epithelial-mesenchymal transition (EMT). CD24 is a surface marker expressed in many tumor types, however, its biological functions and role in cancer progression and treatment resistance remain poorly documented. We have previously shown that loss of CD24 expression in breast cancer cells is associated with radiation resistance, in relationship with the control of oxidative stress. Because ROS are known to mediate the effects of anticancer drugs as well as ionizing radiation, we investigated if CD24 could be defined as an actor of both radiation- and chemo-resistance of breast cancer cells. Using the HMLE breast cancer cell model, we observed that loss of CD24 expression induces stemness properties associated with the acquisition of a hybrid E/M phenotype. The CD24^-/low^ cells were intrinsically more resistant than CD24^+^ cells. The resistance was linked to a lower level of ROS, and CD24 controlled ROS levels through the regulation of mitochondrial functions independently of antioxidant activity. Together, these results suggest a key role of CD24 in de-differentiation process of breast cancer cells, promoting acquisition of therapeutic resistance properties.

## INTRODUCTION

Tumor heterogeneity is a key characteristic of cancer and the presence of cancer stem cells (CSCs) has become increasingly associated with treatment failure and tumor progression/relapse (Marusyk et al., 2012; Pece et al., 2010). CSCs are operationally defined as a subset of cancer cells that *i*) is capable of self-renewal, *ii*) has tumor-initiating ability and *iii*) is resistant to ionizing radiation and chemotherapy. CSCs can be identified by various markers. CSCs from human breast tumors, characterized as having a CD24^-/low^/CD44^+^ phenotype, were first identified for their tumorigenic potential in xeno-transplanted immune-deficient mice (Al-Hajj et al., 2003). Additional markers have been also associated with CSC characteristics and high aldehyde dehydrogenase activity (ALDH^+^), a marker of normal and malignant human mammary stem cells, is a predictor of poor clinical outcome in breast cancer (Ginestier et al., 2007). The subpopulations detected by these markers only partially overlap, indicating that different lineages of CSCs could coexist within the same tumor (Hwang-Verslues et al., 2009).

In addition, much evidence connects epithelial-mesenchymal transition (EMT) with the acquisition of stem cell properties. Many dysregulated pathways in breast CSCs (Notch, hedgehog, Wingless (Wnt), transforming growth factor β (TGFβ) and NFκB), through the activation of several mesenchymal transcription factors (Twist, Snail, Zeb,…), are capable of inducing EMT (Zhang et al., 2021). Most if not all of the mesenchymal cells generated by EMT acquired a CD24^-/low^/CD44^+^ pattern (Mani et al., 2008; Morel et al., 2008; Konge et al., 2018). On the other hand, recent studies clearly showed the existence of Epithelial-like breast CSCs, with a CD24^+^/CD44^low^ phenotype and essentially characterized by high ALDH activity (Liu et al., 2014). Finally, several transition states occurring during EMT have been identified, and a hybrid epithelial/mesenchymal (E/M) phenotype is associated with increased tumor stemness (Pastushenko and Blanpain, 2019; Pasani et al., 2021). In breast cancer, CSCs are endowed with a plasticity that enables them to reverse transition between E and M states, and increased metastatic or tumorigenic potential as well as poor clinical prognosis have been associated with CSCs residing in a hybrid E/M state (Kröger et al., 2019; Pastushenko and Blanpain, 2019).

By definition, breast CSCs are relatively resistant to traditional cancer therapies, including chemotherapy and radiation therapy (Luo et al., 2015), and this resistance has been observed essentially for CD24^-/low^/CD44^+^ M CSCs but also for ALDH^*+*^ E CSCs (Phillips et al., 2006; Li et al., 2008; Kamble et al., 2021; Tanei et al., 2009). Different mechanisms have been proposed to explain the genotoxic stress survival of CSCs (Palomeras et al., 2018), including cell quiescence, increased ability to repair DNA damages, increased antiapoptotic defenses, dysregulation of autophagy, metabolic changes and resistance to reactive oxygen species (ROS), and many of these pathways are mediated by redox imbalance (García-Heredia and Carnero, 2020).

Our group showed previously that high-dose irradiation of breast cancer cell lines leads to the transitory selection of a CD24^-/low^ subpopulation, independently of CD44 expression (Bensimon et al., 2013), and that loss of CD24 expression promotes radiation resistance in relationship with a decreased ROS level (Bensimon et al., 2016). A lower intrinsic level of ROS, associated with radioresistance, is also a characteristic of CD24^-/low^/CD44^+^ M breast CSCs (Diehn et al., 2009; Phillips et al., 2006; Konge et al., 2018).

CD24 is a small cell surface protein molecule and frequently overexpressed in human cancers (Altevogt et al., 2021). CD24 is thought to function as an adhesion molecule, but due to variable glycosylation pattern, it acts as a versatile ligand with diverse physiological functions, making its mechanisms complex to understand. Despite CD24 being a marker strongly associated with EMT in breast cancers, its biological functions and role in cancer progression and treatment resistance remain poorly documented. Moreover, clinical studies associating CD24 expression and breast tumor progression are conflicting, especially due to the poor specificity of CD24 antibodies used in these studies (Kristiansen et al., 2010).

Because ROS are known to mediate the effects of anticancer drugs as well as ionizing radiation, and we previously observed a relationship between CD24 expression and ROS levels, we wondered if CD24 could play a role in the radio- and chemo-resistance of breast cancer cells.

To test this hypothesis, we have used the non-tumorigenic HMLE cell model developed by R. Weinberg’s team (Mani et al., 2008) to study EMT in breast cancer cells, with the main advantage to study E and M cells in the same genetic background. Here, we show that loss of CD24 expression lead to the repositioning of the cells in a hybrid E/M state, displaying increased stemness properties. The CD24^-^ cells are intrinsically more radio- and chemo-resistant than parental cells through the downregulation of ROS level, associated with lower mitochondrial functions. For the first time, these findings provide a functional link between CD24 expression, the hybrid E/M state and therapeutic resistance.

## RESULTS

### CD24 low expression defines a radio- and chemo-resistant subpopulation of breast cancer cells

To investigate the involvement of CD24 in the resistance to γ-irradiation and widely used anticancer drugs, we have studied the CD24 expression in epithelial HMLE cells after high dose irradiation (IR), and after treatment with high concentration of three drugs with different mechanisms of action: 5-Fluorouracil (5-FU), Cisplatin and Paclitaxel. CD24 expression was analyzed by flow cytometry and CD24^-/low^ subpopulation was defined as the 10% of cells expressing the lowest level of fluorescence in untreated cells.

After irradiation, we observed a dose dependent cell death, previously characterized as a delayed apoptotic process (Konge et al., 2018) (Fig. 1A, left). As expected considering our previous results (Bensimon et al., 2013), IR selectively enriches the CD24^-/low^ cells subpopulation (Fig. 1A, right). Similarly, chronic exposure with increased concentrations of anticancer drugs, leads to a dose dependent cell death (Fig. 1B-D, left). As for IR, drug-induced cell death has been strongly associated with late apoptosis in breast cancer (Longley et al., 2003; Tchounwou et al., 2021; Abu Samaan et al., 2019). For the 3 drugs used, a clear increase in the percentage of CD24^-/low^ cells was observed after chronic exposure (Fig. 1B-D, right), again in correlation with cell death. We next investigated if CD24^-/low^ cells are more radio- and chemo-resistant than CD24^+^ cells. These two populations were isolated by flow cytometry (Figure 1E left) and studied separately. Membrane expression level of CD24 observed by FACS was directly correlated to CD24 mRNA expression level in CD24^-/low^ and CD24^+^ cells (Fig. 1E right). After 6 and 10 Gy irradiation, we observed that cell death rate was significantly lower in CD24^-/low^ cells than in CD24^+^ cells (Fig. 1F). In the same way, after treatment with high doses of anticancer drugs, cell death rate was strongly reduced in CD24^-/low^ cells compared with CD24^+^ cells. Therefore, CD24^-/low^ cells have an enhanced ability to survive after irradiation and chronic exposure to chemotherapeutic agents.

**Figure 1:**
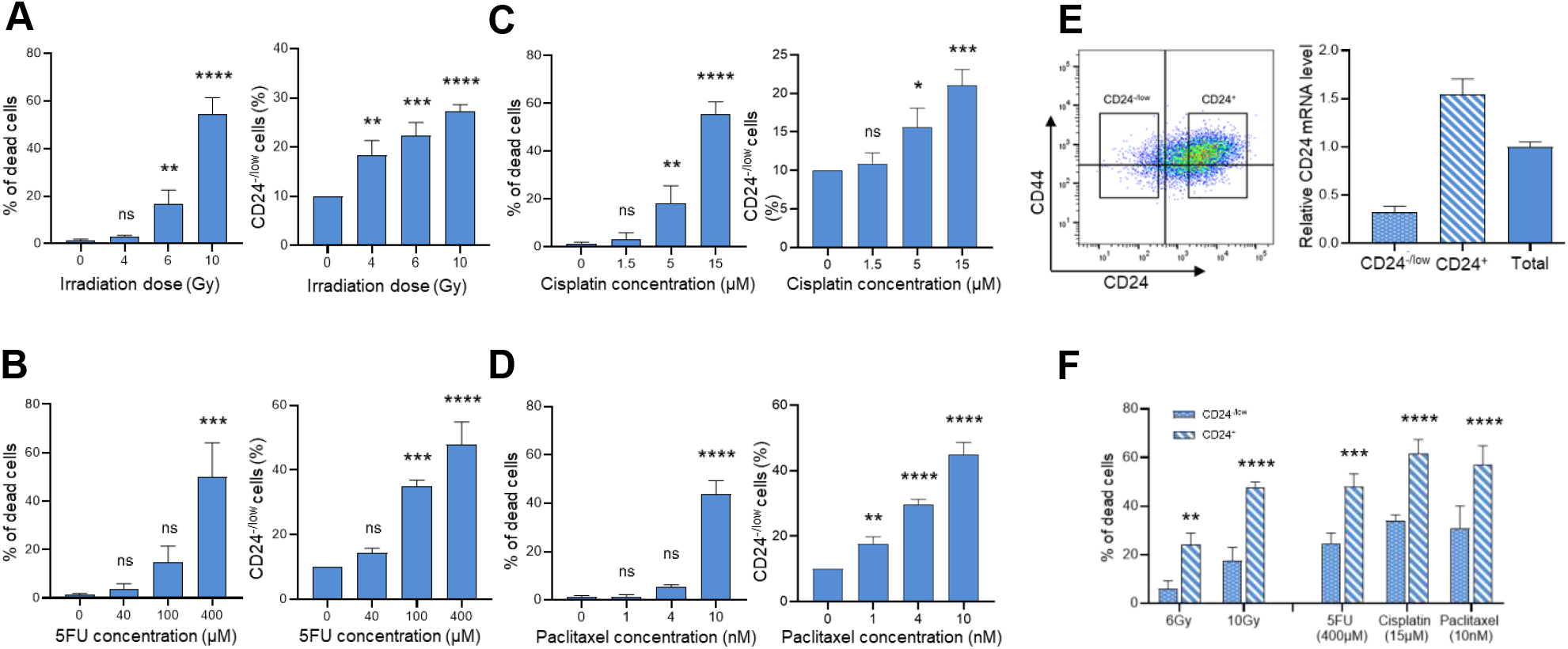
CD24 low expression defines a radio- and chemo-resistant subpopulation of HMLE cells. CD24^-/low^ subpopulation was define as the 10% of cells expressing the lowest level of fluorescence in control cells. (*A*) Percentage of dead cells (*left*) and of CD24^-/low^ cells (*right*) five days after high dose irradiation (4 to 10 Gy) of HMLE cells. (*B, C and D*) Percentage of dead cells (left) and of CD24^-/low^ cells (right) after three days exposure to increasing high concentrations of 5-FU (*B*), Cisplatin (*C*) and Paclitaxel (*D*) of HMLE cells. (*E*) CD24^-/low^ and CD24^+^ HMLE cell subpopulations were FACS sorted after CD24/CD44 staining (*left*) and CD24 expression was analyzed by qRT-PCR (*right*). (*F*) Percentage of CD24^-/low^ and CD24^+^ dead cells five days after 4 and 6 Gy irradiation and after three days exposure to 400μM 5FU, 15μM Cisplatin and 10nM Paclitaxel. For (*A*) to (*F*), results correspond to the mean ± SD of 3 independent experiments. Significant differences between groups were analyzed by ANOVA with multiple comparisons correction test. *P<0.05, **P<0.01, ***P<0.001 and ****P<0.0001.

Taken together, these results indicate that the CD24^-/low^ cells subpopulation is intrinsically more radio- and chemo-resistant than CD24^+^ cells. Hence, irradiation or drug exposure lead to the enrichment of the CD24^-/low^ cells subpopulation in the whole cell culture.

### Loss of CD24 expression does not modify overall epithelial features of HMLE cells, but place cells in a hybrid E/M state

In order to know if CD24 is an actor more than a simple marker of the radio- and chemo-resistance of breast cancer cells, we transfected epithelial (E) HMLE cells with a p-EBV-plasmid expressing an siRNA against CD24 (E_CD24^-^). As control, the parental CD24^+^/CD44^low^ HMLE cells were transfected with p-EBV vector expressing an inefficient shRNA sequence (E_vec). Flow cytometry analyses confirmed the loss of surface expression of CD24 in E_CD24^-^ cells, compared to E cells (Fig. 2A). We also used a purified population of mesenchymal (M) HMLE cells, obtained after FACS sorting of CD24^-/low^/CD44^+^ cells induced after prolonged exposure to TGFβ1 (Konge et al., 2018). In order to discard any possible off-target effects of the siRNA, we generated CD24 complemented E cells (E_CD24^-^c) by transfecting E_CD24^-^ cells with a p-EBV-plasmid coding for the CD24 ORF. A high level of CD24 was observed at the membrane of E_CD24^-^c cells (Fig. 2A). Modulation of CD24 expression levels was further validated by RT-qPCR analysis, showing a down-regulation of about 20 times in E_CD24^-^ cells compared with E or E_vec cells (Supplementary Fig. S1).

**Figure 2:**
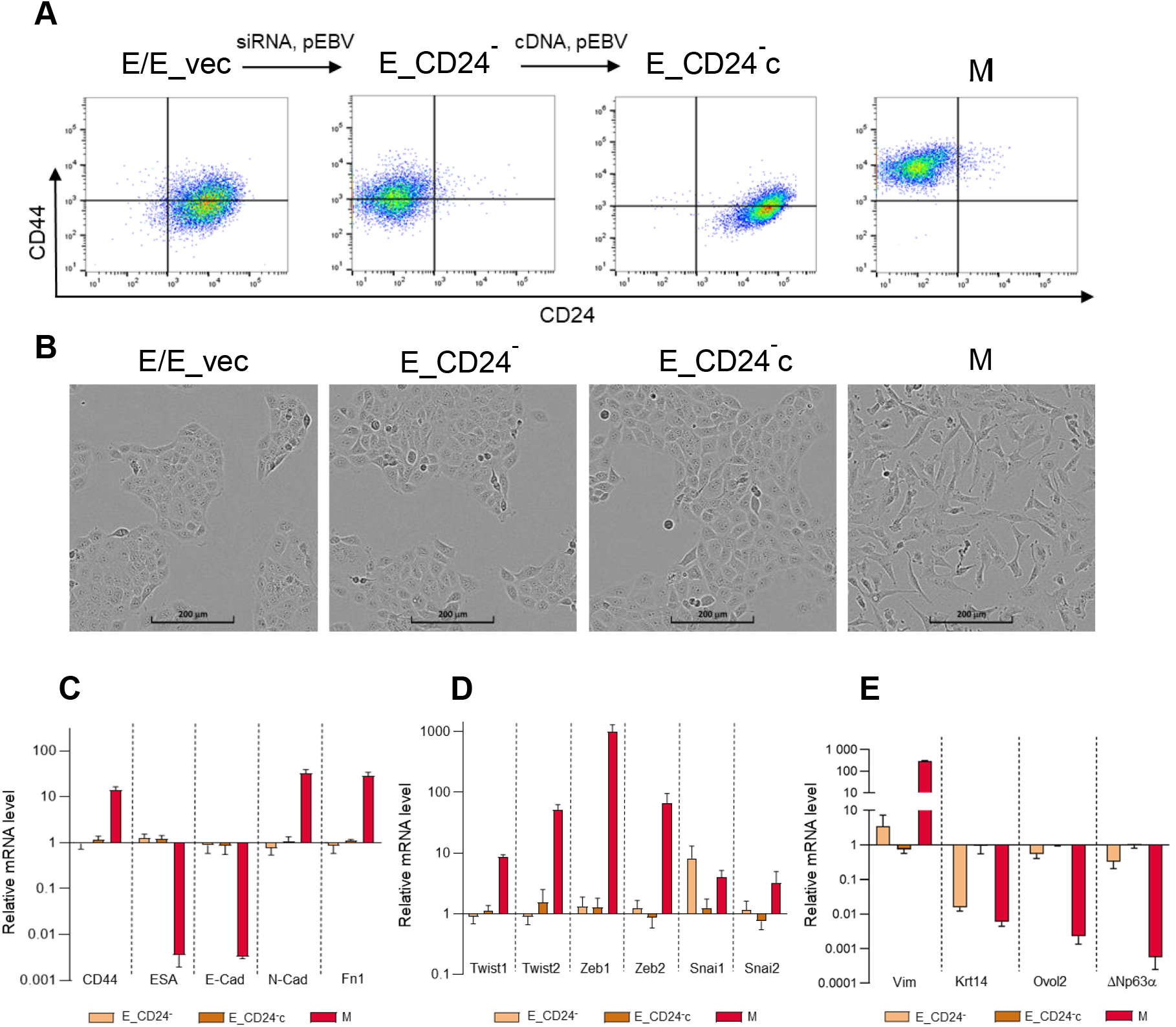
Characterization of E, M and E-transfected HMLE cells. (*A*) Flow cytometry characterization after CD24/CD44 labelling of parental E cells (E), E cells transfected with a control p-EBV vector (E_vec), E cells transfected with a p-EBV-plasmid expressing a CD24 siRNA (E_CD24^-^), E_CD24^-^ cells transfected with a p-EBV-plasmid expressing CD24 mRNA (E_CD24^-^c), and M cells obtained after FACS sorting of CD24^-/low^/CD44^+^ cells induced after prolonged exposure to TGFβ1. (*B*) Phase –contrast images of E / E_vec cells, E_CD24^-^ cells, E_CD24^-^c cells and M cells. (*C, D and E*) Analysis by qRT-PCR of the relative expression of the mRNAs encoding EMT-associated actors (CD44, ESA, E-cadherin, N-cadherin and fibronectin) (*C*), encoding the main transcription factor driving EMT (Twist-1, Twist-2, Snail-1, Snail-2, Zeb-1 and Zeb-2). (*D*), and encoding other actors of EMT (Vimentin, Keratin 14, TP63 and OVOL2) (*E*), in E_CD24-, E_CD24-c and M cells, compared to E cells. Normalization was performed as indicated in materiel and methods. Results were normalized to the expression levels measured in E cells. Each value corresponds to the mean value of 3 independent PCRs performed from 3 independent experiments. Error bars correspond to SEM.

We next examined the influence of CD24 expression on cellular phenotypes classically associated with EMT in breast cancers. E_CD24^-^ cells, as E_CD24c cells, kept the cuboidal-like epithelial morphology of the parental E cells and formed cobblestone cell islands (Fig. 2B). This cobblestone morphology remained different from that observed for the fibroblast-like M cells. To further confirm that silencing of CD24 does not modify the epithelial characteristics of transfected HMLE E cells, the expression of multiple EMT-associated genes was measured by qRT-PCR (Fig. 2C). The results were normalized to expression levels measured in E cells. ESA (Ep-CAM) and E-cadherin, detectable at very low levels in M cells compared to E cells were not affected by the down or up-regulation of CD24. In the same way, CD44, N-cadherin and fibronectin mRNAs expression levels, were markedly increased in M cells but not in transfected E cells. The expression of mRNAs encoding the main transcription factors driving EMT was also studied (Twist-1, Twist-2, Snail-1, Snail-2, Zeb-1 and Zeb-2) (Fig. 2D). Expression of all these mRNAs was increased in M cells but not in E transfected cells with the exception of Snail-1, overexpressed when CD24 expression was reduced. However, expression of other markers suggested that E_CD24^-^ cells were not in a completely epithelial state (Fig. 2E). Keratin 14, a classical epithelial marker in breast cancer (Pastushenko et al., 2018), was down-regulated in E_CD24^-^ cells as in M cells compared to E cells. Vimentin was weakly up regulated and the epithelial transcription factors OVOL2 and ΔNp63α (the predominant p63 isoform in mammary epithelial cells) were weakly under-expressed in E_CD24^-^ cells. Therefore, these results indicated that artificial loss of CD24 expression does not modify overall epithelial characteristics of HMLE.E cells, but induces an intermediate hybrid E/M state.

### Stemness properties and migration potential are associated with the hybrid E/M phenotype of E_CD24^-^ HMLE cells

Stemness properties acquisition during EMT have been associated with the appearance of cells in an intermediate state, often characterized as epithelial-like CSCs displaying an increased ALDH1 activity. We have studied if ALDH1 activity could be related to CD24 expression in HMLE cells (Fig. 3A and Supplementary Fig. S2). About 15-20% of ALDH^+^ cells are present in parental E cells, this subpopulation reached to 40% in E_CD24^-^ cells and re-expression of CD24 (E_CD24^-^c cells) abolished this increase. ALDH^+^ subpopulation is totally absent in CD24^-/low^/CD44^+^ M cells.

**Figure 3:**
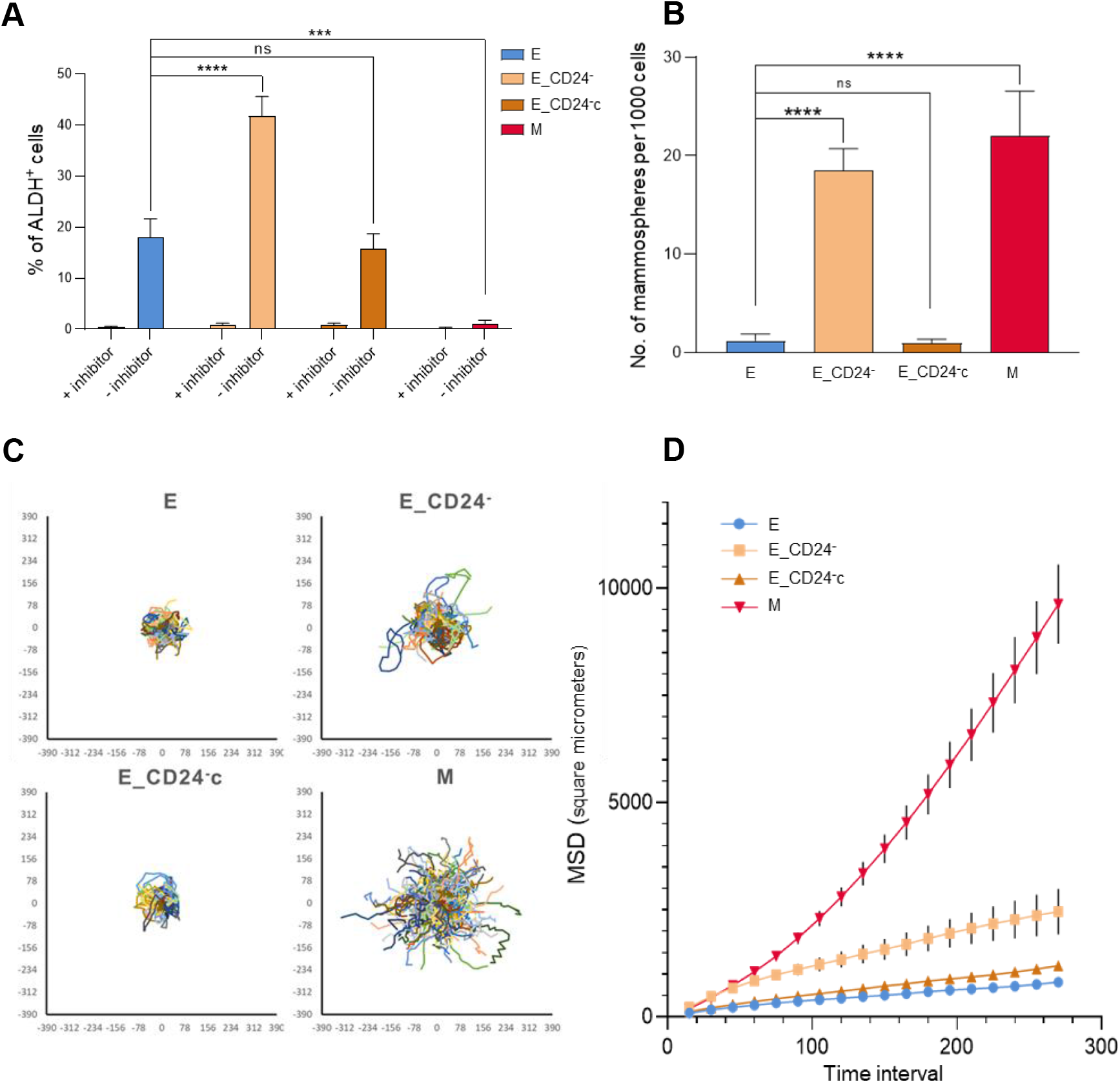
Stemness properties and migration potential of E_CD24-HMLE cells. (*A*) Percentages of the Aldefluor positive subpopulation defined by the Aldefluor assay (see materiel and methods). Results correspond to the mean ± SD of 3 independent experiments. (*B*) Mammosphere formation efficiency. Results correspond to the mean ± SD of 3 independent experiments. For (*A*) and (*B*), significant differences between groups were analyzed by ANOVA with multiple comparisons correction test. ***P<0.001 and ****P<0.0001. (*C and D*) Cell migration potential analysis using the DiPer program (see materiel and methods). (*C*) Cell trajectories visualization representative of three independent experiments. (*D*) Measure of the surface area explored by cells using the MSD analysis. Results correspond to the mean ± SD of 3 independent experiments.

Moreover, another feature of breast CSCs is their potential to form spheres (i.e., mammospheres). As expected, we observed a significant increase in the mammosphere formation efficiency (MFE) of CD24^-/low^/CD44^+^ M cells compared to E cells (Fig. 3B). The MFE was also significantly increased in E_CD24^-^ cells compared to parental E cells, and reexpression of CD24 (E_CD24^-^c cells) abolished this increase.

Finally, as CSCs, as well as hybrid and M tumor cells, have been strongly associated with motility and tumor propagating characteristics (Pastushenko and Blanpain, 2019), we have studied cell migration potential of HMLE cells. Mean square displacements (MSDs) analysis indicated that M cells presents a markedly higher migratory potential than E cells (Fig. 3 C and D). In the same way, E_CD24^-^ cells present an intermediate but significant increase of migratory potential than E cells, and this increase is abolished after reexpression of CD24 (E_CD24^-^c cells).

Altogether, these results indicated that the positioning of E_CD24^-^cells in a hybrid E/M state is associated with the acquisition of stemness properties. Thus, CD24 knock-down could influence stemness properties, leading to ALDH^+^ epithelial-like CSCs rather than CD24^-/low^/CD44^+^ mesenchymal-like CSCs.

### Forced extinction of CD24 expression alone promotes radio- and chemo-resistance of HMLE.E cells

To determine effects of CD24 on radiation and drug sensitivity, HMLE cells were irradiated at 10 Gy or treated with high concentration of 5-FU, Cisplatin and Paclitaxel. Differential delayed cell death was observed between E and M cells during 6 days after irradiation (Fig. 4A). As for M cells, cell death rate after irradiation was significantly reduced in E_CD24^-^ cells compared to parental E cells, and reexpression of CD24 (E_CD24^-^c cells) abolished this resistance. In the same way, the loss of CD24 expression clearly induced a resistance to chronic exposure of anticancer drugs, similar to that observed in M cells (Fig. 4B). When CD24 was re-expressed, E_CD24^-^c cells clearly became drug sensitive again, at the same level as the parental cells. These data indicate that E_CD24^-^ cells have an enhanced ability to survive after irradiation and a chronic treatment with anticancer drugs.

**Figure 4:**
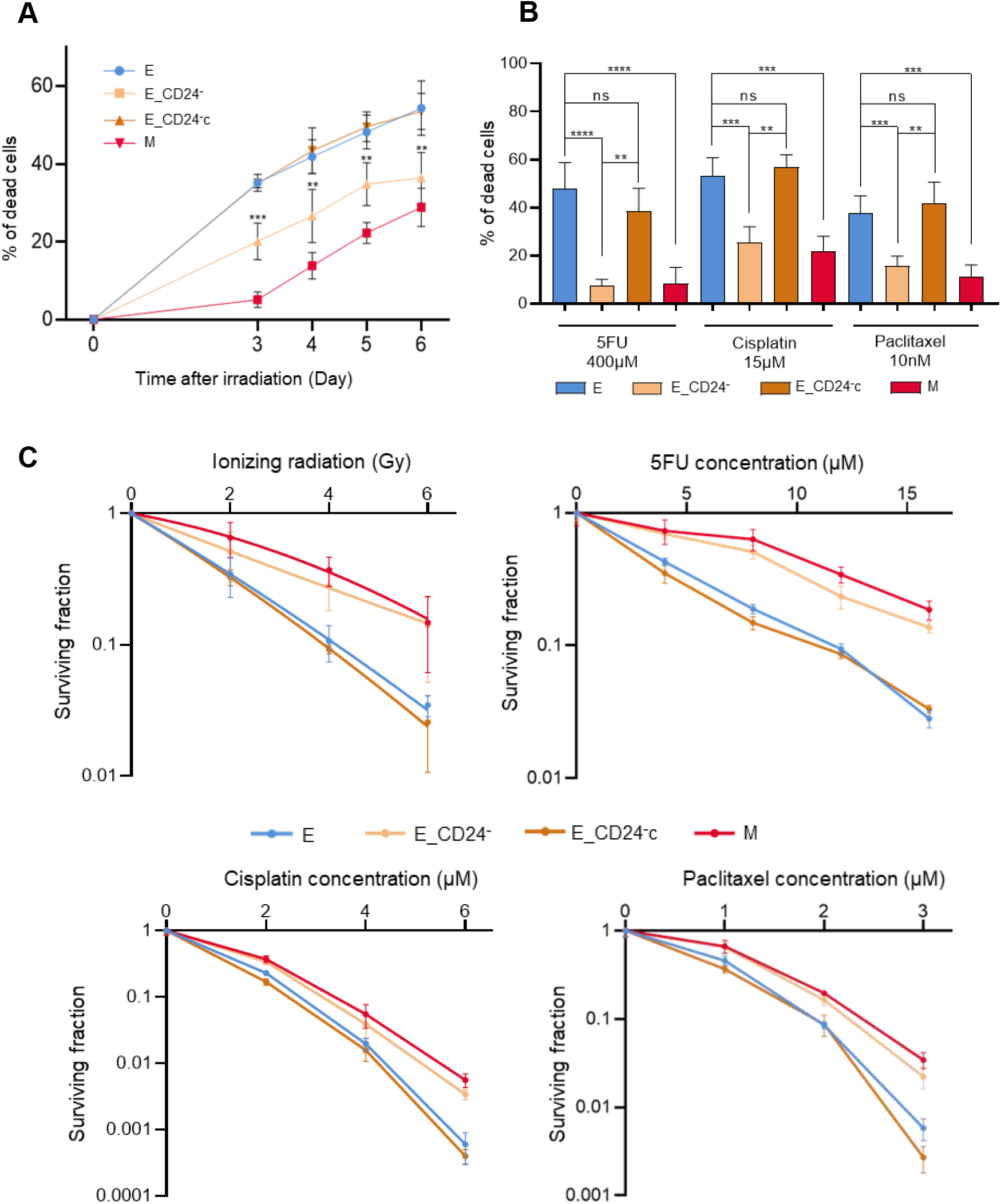
Reduced CD24 expression enhances radio- and chemo-resistance in HMLE.E cells. (*A*) Time course of cell death of 10 Gy-irradiated cells. Results correspond to the mean ± SD of 3 independent experiments. **P<0.01, ***P<0.001. (*B*) Percentage of dead cells after three days exposure to 400μM 5FU, 15μM Cisplatin and 10nM Paclitaxel. Results correspond to the mean ± SD of 4 to 12 independent experiments. For (*A*) and (*B*), significant differences between groups were analyzed by Mann-Whitney tests. **P<0.01, ***P<0.001 and ****P<0.0001. (*C*) Clonogenic cell survival curves after: 2 to 6 Gy irradiation, 4 to 16 μM exposition to 5FU, 2 to 6 μM exposition to Cisplatin and 1 to 3 μM exposition to Paclitaxel.

To ensure that these results are not specific to our HMLE cells model, two other epithelial breast cell lines, MCF7 and T47D, were also transfected with the p-EBV-plasmid expressing a CD24 siRNA, in order to obtain two cell lines with a low CD24 expression (MCF7_CD24^-^ and T47D_CD24^-^ cells). As for HMLE cells, forced extinction of CD24 expression alone promotes resistance of MCF7_CD24^-^ and T47D_CD24^-^ cells against chronic exposure to high dose of 5-FU and Cisplatin (Supplementary Fig. S3).

We next investigated whether differences in the cell death rate after irradiation or drug treatment between CD24^-^ and CD24^+^ cells had an impact on long-term survival, as measured by the clonogenic cell survival assay after treatment. After irradiation (Fig. 4C), surviving fractions at 2, 4 and 6 Gy were significantly higher in E_CD24^-^ than in control E/E_vec cells, and similar to that observed in M cells. Complementation of CD24 (E_CD24^-^c cells) led to a decrease of cloning efficiency, like in parental E cells. After drug treatments, similar results were observed: surviving fractions for all doses of drug tested were higher in E_CD24^-^ and M cells than in parental E cells and E_CD24^-^ cells. These results indicate that E_CD24^-^ cells, as M cells, generate a larger progeny than CD24^+^ E cells.

Taken together, these data provide evidence that CD24 protein controls the response to radiation and chemotherapeutic drugs.

### Reduced CD24 expression decreases intracellular ROS concentration

As ROS production is reported to be an essential inducer of apoptosis, we thus studied whether changes in ROS level could be related to radio- and chemo-sensitivity observed in HMLE cells. First, we measured the intracellular concentrations of ROS using dihydroethidium (DHE) staining. Flow cytometry analysis indicated that M cells contained significantly lower concentration of ROS than E/E_vec cells (Fig. 5A, left), in agreement with our previous results (Bensimon et al., 2013) and the recurrent observation of low ROS levels in CD24^-/low^/CD44^+^ breast cancer cells (Diehn et al., 2009; Phillips et al., 2006). E_CD24^-^ cells also display a lower DHE staining than E control cells and the reexpression of CD24 abolished this decreased staining. Because mitochondria is the major source of ROS in cancer cells, a similar study was performed using Mitosox-Red, a selective probe for mitochondrial superoxide and identical results were observed (Fig. 5B, left).

**Figure 5:**
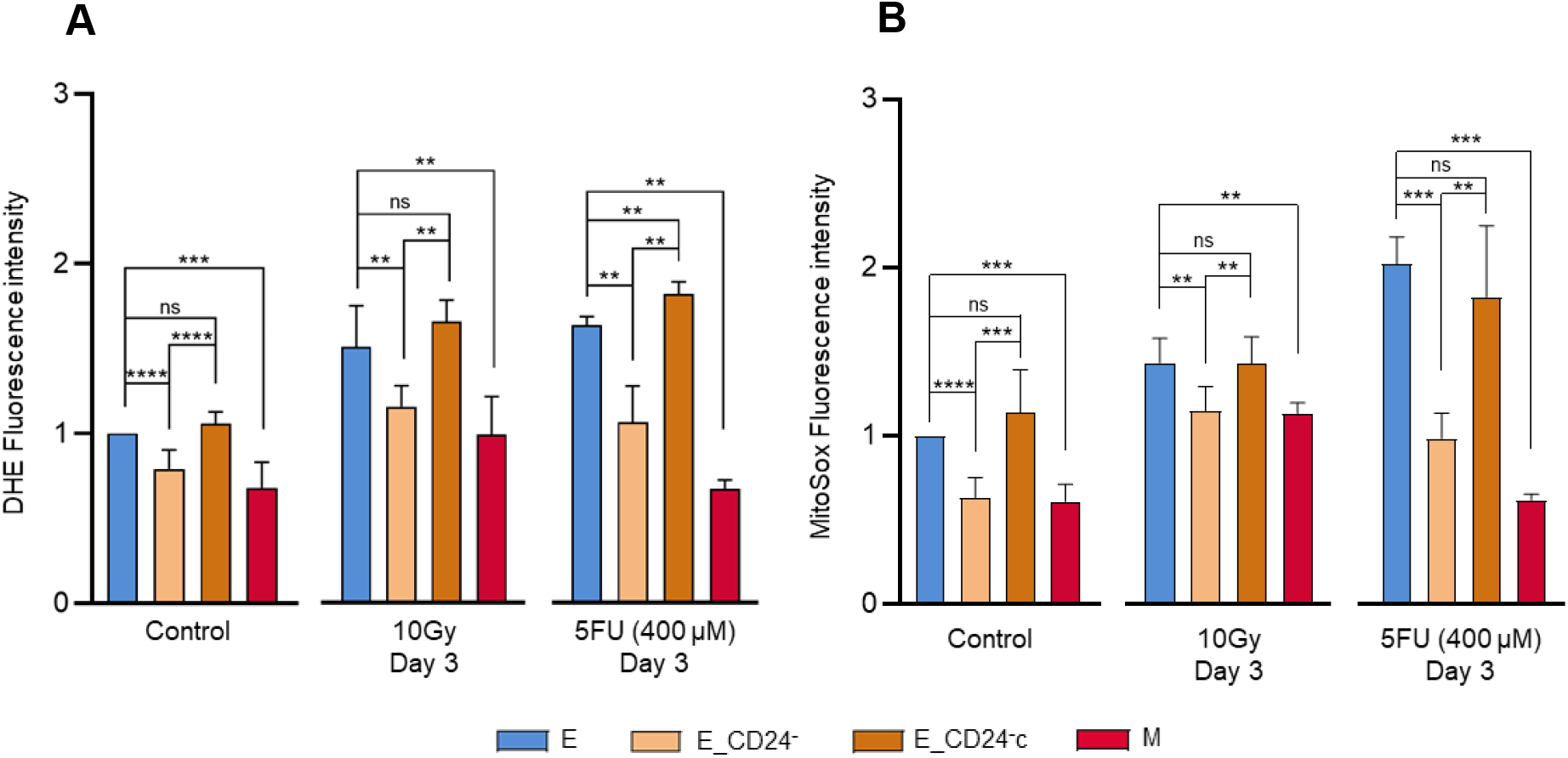
CD24 down regulation decreases total as well as mitochondrial ROS level. (*A*) Cellular ROS level was assessed by DHE probe and (*B*) mitochondrial ROS level was assessed by Mitosox-Red probe. For (*A*) and (*B*), ROS level were studied in untreated cells (left), three days after 10 Gy irradiation (middle) and three days after exposure to 400μM 5FU (right). Results correspond to the mean ± SD of 4 to 10 independent experiments. Significant differences between groups were analyzed by Mann-Whitney tests. **P<0.01, ***P<0.001 and ****P<0.0001.

We next investigated the impact of CD24 expression on ROS levels 3 days after irradiation or 5FU treatment. ROS level increased in all cell lines tested (Fig. 5 A and B, right), but a lower concentration of ROS was maintained in E_CD24^-^ cells (as for M cells) compared to parental E or E_CD24^-^c cells. Similar results were obtained with the two probes used: DHE and Mitosox-Red.

Taken together, these data indicate that CD24 down regulation leads to decreased basal level of total as well as mitochondrial ROS. After irradiation or drug treatment, we observed an increase in ROS level, but it remains lower in E_CD24^-^ cells than in parental E cells, consistent with the rate of cell death observed in the different cell lines. Therefore, CD24 expression could impact radio- and chemoresistance through the control of ROS level.

### CD24 controls ROS level through the regulation of mitochondrial functions independently of antioxidant activity

A lower ROS level is commonly ascribed to the CSC phenotype in breast tumors, and has been related to enhanced ROS scavengers and/or decreased mitochondrial mass (Peiris-Pagès et al., 2016). First, we determined whether ROS modulation could be related to differential regulation of oxidative stress-related genes. We previously showed that HMLE.M cells presented a higher antioxidant activity than HMLE.E cells, and characterized a set of genes involved in ROS metabolism (Konge et al., 2018). Thus, we studied by RT-qPCR the expression of four key genes in this model: SOD2, HMOX1, GSR, TXNRD1 (Fig. 6A). The expression of all these genes is increased in M cells, but not modified in transfected E cells. So, the lower ROS level observed in E_CD24^-^ cells is not related to high intracellular levels of radical scavengers.

**Figure 6:**
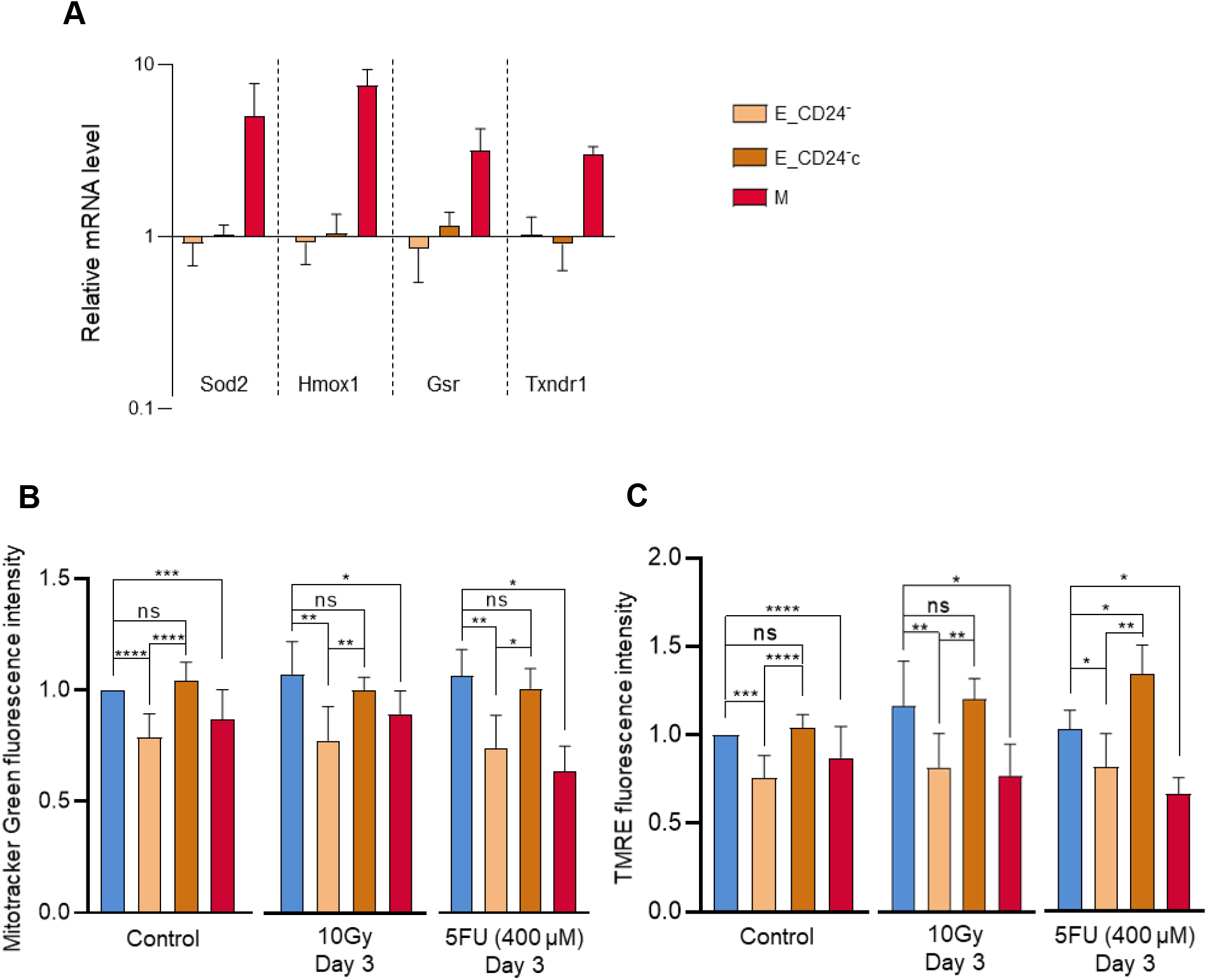
CD24 down regulation have no impact on ROS scavenger but decreases mitochondrial mass and mitochondrial membrane potential. (*A*) Analysis by qRT-PCR of the relative expression of the mRNAs encoding stress-related actors involved in ROS metabolism in E_CD24-, E_CD24-c and M cells, compared to E cells. Normalization was performed as indicated in materiel and methods. For each gene, expression in E cells was normalized to 1 and the ratio of relative mRNA level of E cells to E_CD24-, E_CD24-c and M cells was presented. Each value corresponds to the mean value of 3 independent PCRs performed from 3 independent experiments. Error bars correspond to SEM. (B) Mitochondrial mass was assessed by Mitotracker Green probe and (**C**) mitochondrial membrane potential was assessed using tetramethylrhodamine, ethyl ester (TMRE) staining. For (*B*) and (*C*), mitochondrial mass and mitochondrial membrane potential were studied in untreated cells (left), three days after 10 Gy irradiation (middle) and three days after exposure to 400μM 5FU (right). Results correspond to the mean ± SD of 4 to 10 independent experiments. Significant differences between groups were analyzed by Mann-Whitney tests. *P<0.05, **P<0.01, ***P<0.001 and ****P<0.0001.

We next studied if modulation of CD24 expression have an impact on mitochondrial functions. Mitochondrial mass was analyzed using Mitotracker Green (MTG), a mitochondrial selective probe, and mitochondrial membrane potential was quantified using tetramethylrhodamine, ethyl ester (TMRE) staining. Flow cytometry analysis indicated that mitochondrial mass was significantly lower in M cells and in E_CD24^-^ cells than in parental E and E_CD24^-^c cells (Fig. 6B, left). In the same way, mitochondrial membrane potential decreased in M cells and in E_CD24^-^ cells compared to E cells, and a return to the initial level was showed in E_CD24^-^c cells (Fig. 6C, left). The TMRE/MTG ratio is identical for all the cell lines analyzed (Supplementary Fig. S4), indicating that variation of mitochondrial membrane potential observed in the cell lines is the direct consequence of variation of mitochondrial mass. Then, similar studies were carried out after irradiation or drug treatment (Fig. 6 B and C, right). As for untreated cells, mitochondrial mass and mitochondrial membrane potential are lower in M cells and in E_CD24^-^cells, compared to parental E cells and E_CD24^-^c cells.

Taken together, these data indicated that the lower ROS levels observed in CD24^-/low^ cells is related to a lower mitochondrial membrane potential associated to a lower mitochondrial mass, independently of the intrinsic antioxidant activity. So, a markedly reduced level of mitochondrial ROS production appears to be a key event in the radio- and chemo-resistance controlled by CD24 expression.

## DISCUSSION

Our data provide evidences that intrinsic radio- and chemo-resistance of breast cancer cells is directly linked to the membrane expression level of CD24, and that CD24 can be considered not only as a marker but also as an actor of this resistance. It was shown that CD24 expression presents a dynamic regulation (Meyer et al., 2009), its expression being reversible and under epigenetic control. Expression of CD24 is heterogenous in our HMLE cell model (Fig. 1E), as it is frequently observed in breast cancer cell lines and in tumor tissues (Ricardo et al., 2011).

In the literature, radio- and chemo-resistance of breast cancer cells have been highlighted using different protocols of irradiation or different classes of chemotherapeutic drugs. In our study, we chose to use high doses of irradiation (4 to 10 Gy) and chronic exposure with a high concentration of drugs from three categories of antitumor agents: 5FU is an antimetabolites drug, Cisplatin an alkylating agent and Paclitaxel a mitotic inhibitor. In parental HMLE.E cells (Fig. 1), we observed a dose dependent cell death, after irradiation as well as after drug treatment, and this cell death has been mainly associated with late apoptosis (Konge et al., 2018; Luce et al., 2009; Longley et al., 2003; Tchounwou et al., 2021; Abu Samaan et al., 2019). In all experiments, cell death is concomitant with a clear increase in the percentage of CD24^-/low^ surviving cells. This enrichment in CD24^-/low^ cells could be the consequence of a selection (Bensimon et al., 2013), but also related to the induction of an active EMT program (Pinto et al., 2013). When we compared the resistance of FACS-sorted CD24^-/low^ cells and CD24^+^ cells (Fig. 1 *E* and *F*), we observed that CD24^-/low^ cells had a higher ability to survive than CD24^+^ cells after irradiation or antitumor drugs treatment. Altogether, our results associate CD24^-/low^ expression with treatment resistance, independently of the mechanism of CD24^-/low^ cells enrichment.

Failure of conventional treatments has been commonly associated with CSCs survival (Luo et al., 2015), suggesting that E_CD24^-^ cells have acquired stemness properties. In the last 20 years, numerous studies have associated loss of CD24 expression with EMT. The widely used CD24^-/low^/CD44^+^ marker allows characterizing M cells, enriched in CSC properties compared to CD24^+^/CD44^low^ E cells (Al-Hajj et al., 2003; Mani et al., 2008; Konge et al., 2018). However, in our study, E_CD24^-^ cells retained the main characteristics of E cells: cuboidal-like epithelial morphology and formation of cobblestone cell islands. Furthermore, the expressions of the main EMT-associated genes (ESA, E-cadherin, CD44, N-cadherin and fibronectin) and 5 out of 6 of the transcription factors driving EMT were not modified (Fig. 2). Therefore, treatment resistance observed in E_CD24^-^ cells does not seem to be associated with acquisition of M characteristics.

The development of biomarkers to identify breast CSCs made it possible to show that cells with high ALDH activity present also stemness properties and radiation- and chemo-resistance properties have been also associated with ALDH^+^ breast cancer cells (Phillips et al., 2006; Li et al., 2008; Kamble et al., 2021; Tanei et al., 2009). Loss of CD24 expression may promote the appearance/increase of stemness characteristics allowing the genotoxic stress survival of E_CD24^-^ cells, radio- and chemoresistance of E_CD24^-^ cells being correlated to ALDH activity. Moreover, in our study, the idea of stem cell features associated with E_CD24^-^ cells is reinforced by the increase potential to form mammospheres and the increase migratory potential observed for these cells (Fig. *3*B-D).

In the literature, the initial view that only a “full” EMT was associated with increased stemness was challenged by later studies demonstrating the existence of Epithelial-like breast CSCs (Liu et al., 2014). These E CSCs with a CD24^+^/CD44^low^ phenotype are essentially characterized by high ALDH activity. Finally, a more nuanced recent view highlights that epithelial-mesenchymal plasticity is a spectrum of transitory cell states, and tumor cells with the highest stem cell capabilities reside in a hybrid E/M state, associated with an increased tumor propagating potential (Kröger et al., 2019; Bierie et al., 2017; Pastushenko et al., 2018; Pasani et al., 2021).

In our study, if overall epithelial characteristics of HMLE cells are maintained, deregulation of other genes associated with transition states occurring during EMT is also observed: Snail-1, Vimentin, Keratin 14, p63 and OVOL2 (Figure 2). The strong down-regulation of Keratin 14 expression (E marker) and the low but significant overexpression of Vimentin (M marker), suggest that E_CD24^-^ cells have acquired few M characteristics (Fig. 2E). Among the main EMT transcription factors, only Snail1 is strongly overexpressed in E_CD24^-^ cells (Fig. 2B). Snail1 expression has been implicated in reprogramming process of somatic cells to pluripotency (Unternaehrer et al., 2014; Gingold et al., 2014), and Snail1 also maintains stem-like properties, chemoresistance and ALDH activity in mouse breast cancer cells (Ma et al., 2017). Importantly, Krôger et al. have shown that, both *in vitro* and *in vivo*, breast cancer cells can reside stably and with low cell plasticity in a highly tumorigenic hybrid E/M state, which is driven mainly by Snail1 and canonical Wnt signaling (Kröger et al., 2019). The epithelial transcription factors p63 and OVOL2 were weakly but significantly down expressed in E_CD24^-^ cells but at a lower level than in M cells (Fig. 2E). Repression of p63 has been implicated in EMT induction in mammary epithelial cells (Yoh et al., 2016). OVOL2 is a key regulator of epithelial-mesenchymal plasticity as well as CSCs (Pasani et al., 2021). Expression of OVOL2 induce MET, antagonizes TGF-β signaling and has been implicated in positioning mammary epithelial cells in an intermediate E/M state (Roca et al., 2013; Wu et al., 2017; Hong et al., 2015). Finally, E_CD24^-^ cells present a significant increase of migratory potential, which can be strongly associated with the increased tumor propagating potential observed in breast cancer cells in a hybrid E/M state (Pastushenko and Blanpain, 2019). Altogether, these data reinforced the idea that the hybrid E/M state of E_CD24^-^ cells lead to the acquisition of stemness, and the maintenance of stem cell properties is independent of phenotypic plasticity.

The cellular model used in our study has the advantage of using cells with the same genetic background and the plasticity between E and M allows the characterization of the alterations induced by the regulation of CD24 expression levels in E cells and a fine mapping in the E/M hybrid state. This model not only allows us to generalize our previous observations (Bensimon et al., 2016) but also to go much further in the understanding of the sequence of events and to measure the direct consequences of the down-regulation of CD24.

The intrinsic radio- and chemo-resistance of CSCs have been associated with many mechanisms, often mediated by redox imbalance, and ROS control appears to be a key mechanism in this resistance (García-Heredia and Carnero, 2020). Moreover, ROS production is reported to be an essential regulator or inducer for apoptosis in cancer cells, and an important increase of intracellular ROS level is known to mediate cell death induced by ionizing radiation (Riley, 1994) as well as anticancer drugs (Mohiuddin and Kasahara, 2021; Mosca et al., 2021; Pan et al., 2013). Increased ROS scavengers have been extensively associated to the lower ROS level observed in CSCs (Diehn et al., 2009; Konge et al., 2018), but few papers also report that EMT CSCs showed decreased mitochondrial mass and membrane potential, consumed far less oxygen per cell, and produced markedly reduced levels of ROS (Gammon et al., 2013). Our results demonstrate that CD24 down regulation leads to decreased basal level of total as well as mitochondrial ROS, through the regulation of mitochondrial functions. A link between mitochondrial repression and metabolic reprogramming has been proposed, and the mitochondrial lower levels and activity can reflect the metabolic switch that has been reported during EMT (Jia et al., 2021; Lee et al., 2019). Moreover, it appears that Snail plays a central role in this process. Altogether, our results show a direct relationship between CD24 expression and therapeutic resistance. In breast cancer cells culture, CD24 expression is heterogeneous, and CD24^-/low^ subpopulation is clearly selected after radiation and drug treatment. Artificial modulation of CD24 expression shows that radio- and chemo-sensitivity are directly under the control of CD24, and that loss of CD24 expression in E HMLE cells increase the presence of ALDH^+^ Epithelial-like CSCs, in a hybrid E/M state. As for CD24^-/low^/CD44^+^ CSCs, the resistance properties are strongly associated to a low ROS level, but for the two subtypes of CSCs the mechanisms leading to a decrease ROS level are different. For ALDH^+^ E_CD24^-^ cells, we observed a decrease of mitochondrial mass and mitochondrial membrane potential, while for CD24^-/low^/CD44^+^ M cells, decrease ROS level is under the control of both enhanced ROS scavenger and decrease of mitochondrial mass and mitochondrial membrane potential. These results can be summarized in a model presented in Fig. 7.

**Figure 7:**
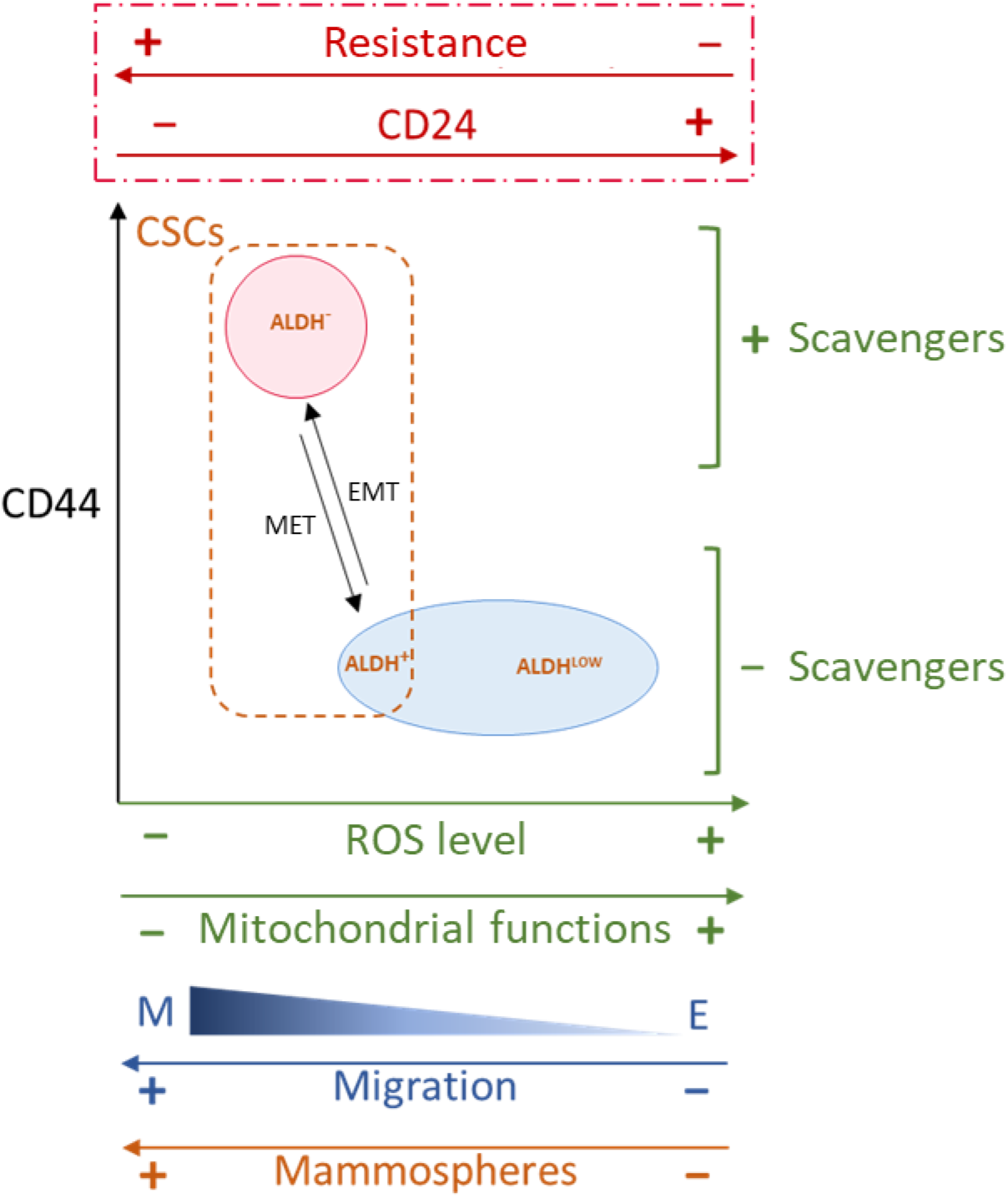
Proposed model of the relationships between CD24 expression and EMT, stemness properties, ROS level, ROS scavengers and mitochondrial functions.

In the last decade, few papers have studied the role of CD24 in human tumors, and even if different functions of CD24 have been proposed, its role in cancer progression and treatment resistance remain poorly documented, and the signaling downstream of CD24 has not been clearly elucidated yet (Altevogt et al., 2021). In breast tumors, high CD24 expression is frequently associated with a terminally differentiated, luminal phenotype, while most of the basal-like tumors are classified as CD24^-/low^. In parallel, studies have shown that CD44/CD24 and ALDH1 are expressed differentially in different subtypes of breast cancers. The ALDH^+^ cells are more common in HER2-overexpressing and basal/epithelial breast cancer, while the CD24^-/low^/CD44^+^ phenotype is more associated with basal like breast cancer (Li et al., 2017; Ricardo et al., 2011). Moreover, only a fraction of CD24^-/low^/CD44^+^ breast cancer cells are ALDH^+^, these cells being tumorigenic (Ginestier et al., 2007; Liu et al., 2014). Our work strongly suggests that CD24 expression level may be a key factor in the transition between different subtypes of breast CSCs and that loss of CD24 expression is connected to a de-differentiation process. De-differentiation is classically related to invasion potential and mestastasis, and is also sufficient to promote resistance to a wide spectrum of chemotherapy drugs and radiation exposure (Gupta et al., 2019).

## ACKNOWLEDGMENTS

This work was funded by Electricité de France (EDF) and by the Transverse Division n. 4 (Radiobiology) of the French Alternative Energies and Atomic Energy Commission (Segment n. 4 Radiobiologie)

## AUTHOR CONTRIBUTIONS

I.B., A.C. and J.L. designed research; I.B., C.L., N.D., T.K., E.B., C.S. and J.L. performed research; T.K. and F.B. contributed new reagents/analytic tools; I.B., S.C., A.C. and J.L. analyzed data; I.B., E.B., F.B., A.C., and J.L. proofread the paper and I.B., A.C. and J.L. wrote the paper.

## DECLARATION OF INTERESTS

The authors declare no competing interests.

## Materials and methods

### Cell culture

All of the cell lines used for this study were cultivated in Heraeus Thermo Scientific™ BBD6220 incubator at 37°C in a humidified atmosphere of 5% CO_2_ and 95% air. Presence of mycoplasma were regularly tested with the MycoAlert™ Mycoplasma detection kit from Lonza. Tumor breast cancer cell lines T47D and MCF7 were from the American Type Culture Collection (Rockville, MD) and Human Mammary Epithelial HMLE cell line was kindly provided by RA Weinberg (Cambridge, MA, USA). All cell lines were grown in adherent conditions but in different cell culture media always supplemented with 10% (v/v) heat-inactivated fetal bovine serum (Sigma-Aldrich, Saint-Louis, MO – USA) and 1 mM antibiotic-antimycotic (Invitrogen, Carlsbad, CA – USA). For T47D and MCF7, Dulbecco’s Modified Eagle’s Medium (DMEM), high glucose, GlutaMAX™ Supplement, pyruvate (Gibco) was used therefore HMLE cells were cultured in DMEM/F-12 Nutrient Mixture, GlutaMAX™ Supplement (Gibco) extemporaneously completed by 10 ng/ml human epidermal growth factor (EGF) (Sigma), 0.5 μg/ml hydrocortisone (Sigma) and 10 μg/ml insulin (Sigma). Medium for transfected cell lines was supplemented with 0.4 μg/mL puromycin (Gibco) and 16 μg/mL blasticidin (Gibco) during the transfectant selection phase, and 0.2 μg/mL puromycin and 8 μg/mL blasticidn in routine cell culture. HMLE.M cells were induced after prolonged treatment with 2.5 ng/ml recombinant TGFβ1 (Life Technologies, Cergy-Pontoise, France). Cell proliferation and survival analyses were performed in 3 or more independent experiments, by scoring cells with a TC20 automated cell counter (BIO-RAD). Discrimination between viable and dead cells was performed by trypan blue exclusion. Mammosphere formation assay were performed as described in Lombardo et al. (Lombardo et al., 2015) (*SI Appendix, Supplementary Materials and Methods*).

### Chemical, reagents and antibodies

5-Fluorouracil (5FU), Cisplatin and Paclitaxel (Taxol) were from Sigma-Aldrich (Saint-Louis, MO, USA). 5FU and Paclitaxel were diluted in dimethylsulfoxid (DMSO) solution at concentration stock of 400mM and 1mM respectively. Cisplatin was thinned with sodium chloride at concentration stock of 1Mm. Each drug was used at different concentrations depend on the experiments. Antibodies against CD24 (clone ML5), CD44 (clone G44-26), and isotypic controls were from BD Biosciences (San Jose, CA, USA).

### Plasmids and transfection

To stably knock down the expression of CD24 in MCF-7, T47D and HMLE cells, we used replicative short hairpin (shRNA)-expressing vectors (puromycin-resistant pEBV-siRNA), which impose a very strong and stable gene silencing in human cells, even after several months in culture. siRNA design and cloning in pEBVsiRNA vectors but also establishment of stable knockdown and control clones were carried out as previously described (Biard et al., 2005). To design shRNA sequences, we adopted the DSIR program developed by Vert *et al*. (Vert et al., 2006). Control cells carried a plasmid expressing an inefficient shRNA sequence called E_vec.

To complement CD24 expression into HMLE cells already depleted for CD24, open reading frame (ORF) of human CD24 was amplified from an I.M.A.G.E. clone (ID_5591617, Thermo Scientist) with the following primers: 5’-ATGGGCAGAGCAATGGTGGCCAGGCTC-3’ and 5’-TTAAGAGTAGAGATGCAGAAGAGAGAG-3’ and introduced into a blasticidin-resistant pEBV plasmid downstream of a CAG promoter.

After one day of seeding into 6-well plate (TPP, Dutscher), cells were transfected with jetPRIME^®^ reagent (Polyplus, Illkrich, France) according to the manufacturer’s recommendations. After 24-h incubation, cells were trypsinized and seeded in culture medium supplemented with puromycin alone or with blasticidin. Experiments were performed either on the whole transfected population or on selected clones.

### Irradiation

For all experiments except clonogenic assays, cells were plated at least 24h prior to irradiation. On day 0, γ-irradiations of cells were performed on a GSR D1 irradiator (Gamma Medical Service GmbH) (*SI Appendix, Supplementary Materials and Methods*). Studied cells were irradiated at 2, 4, 6 and 10 Gy and control cells were submitted to sham irradiation.

### ALDEFLUOR assay

The ALDEFLUOR kit (StemCell Technologies, Durham, NC, USA) was used to analyze the population with high ALDH enzymatic activity according to the manufacturer’s intructions. The cells were incubated in the ALDEFLUOR assay buffer containing ALDH substrate (BAAA, 1 μmol/l per 1×10^6^ cells) and incubated during 40 minutes at 37°C. As negative control, for each sample of cells, an aliquot was treated with 50mmol/L diethylaminobenzaldehyde (DEAB), a specific ALDH inhibitor.

### Clonogenic assays

Sub-confluent cells were trypsinized using TrypLE express solution (Thermofisher). Living cells were counted using an automated cell counter (TC20, Biorad) considering trypan blue exclusion. To this point, the protocol differs slightly depending on cellular treatment. Irradiation: cells were immediately irradiated in suspension for the time required to generate a dose curve of 0, 2, 4 and 6 Gy and colony-forming assays were performed immediately after irradiation by plating cells in 60 mm diameter Petri dishes, in triplicate.

#### Drug treatment

after trypsinization and counting, cells were plated in 60 mm diameter Petri dishes, in triplicate. Six hours after plating, drug treatment was performed for three days, and then cells were washed and incubated with fresh medium.

Number of cells seeded increase with radiation dose or drug concentration, but was identical for each cell line tested. After 7 days, cells were fixed for 30 minutes in 4% paraformaldehyde, washed and stained for at least two hours in methylene blue/30% methanol. Colonies containing more than 50 cells were counted. The surviving fraction at each radiation dose was normalized to that of the nonirradiated sample, and points were fitted using an exponential tendency curve. At least three independent experiments were performed.

### Cell staining

Cancer stem cell marker labeling and analysis were performed as described in Bensimon et al.(Bensimon et al., 2013, 2016) Intracellular concentrations of ROS prooxidants were determined using dihydroethidium (DHE) probe (Invitrogen™). Adherent cells were incubated with 10μM DHE for 30 min at 37°C, then washed with PBS, trypsinized and immediately analyzed by flow cytometry. Intracellular concentrations of mitochondrial ROS were determined using MitoSOX™ Red Mitochondrial Superoxide Indicator (Invitrogen™). Adherent cells were incubated with 5μM MitoSOX Red for 10 min at 37°C, then washed with PBS, trypsinized and immediately analyzed by flow cytometry. Mitochondrial mass was analyzed using MitoTracker™ Green (MTG) probe (Invitrogen™). After soft trypsinization cells were loaded with 200 nM MTG and incubated for 20 min at 37°C, then immediately analyzed by flow cytometry. Mitochondrial membrane potential was quantified using tetramethylrhodamine, ethyl ester, perchlorate (TMRE) probe (Invitrogen™). After soft trypsinization cells were loaded with 10 nM TMRE and incubated for 20 min at 37°C, then immediately analyzed by flow cytometry.

### Flow cytometry

Cells were analyzed on a SORP LSR-II analyzer (Configuration: 488 nm, 561 nm, 405 nm, 355 nm, and 635 nm) or on a BD FACSCalibur (Configuration: 488 nm and 635 nm) (BD Biosciences, San Jose, CA). Data were analyzed with FlowJo v10.7.1 (Tree Star). Cells were sorted on a BD Influx sorter (BD Biosciences) (Configuration: 488 nm, 561 nm, 405 nm, 355 nm, and 635 nm).

### RNA extraction and quantitative real-time PCR

Total RNA was extracted from frozen cell pellets with the « total RNA purification kit » (Norgen Biotek Corp), according to the manufacturer’s intructions. DNAse treatment was realized using the TURBO DNA-free Kit (Invitrogen™ Ambion™), according to the manufacturer’s intructions. cDNA synthesis was performed with the SuperScript™ VILO™ cDNA Synthesis Kit (Invitrogen™) according to the manufacturer’s intructions. RT-PCR was performed with an ABI Prism 7300 detection apparatus (Applied Biosystems, Courtaboeuf, France) using the Taqman Universal Master Mix according to the manufacturer’s recommendations. The Ct value was determined with the Sequence Detection System software. All the primers were from Applied Biosystems (*SI Appendix, Supplementary Materials and Methods*). Levels of gene expression were determined using GENORM software and normalized using GAPDH and RPLPO.

### Cell migration analysis

Cells were plated on 24 well glass bottom dishes at 4000 cells/well density. Live-cell imaging was performed through a Plan APO 20x DIC objective (NA: 0.7) on a Nikon A1R confocal laser scanning microscope system attached to an inverted ECLIPSE Ti (Nikon Corporation, Tokyo, Japan) thermostated at 37°C under a 5% CO2 atmosphere. Mosaic images were recorded every 15 minutes during 5 hours. Tracking of individual cells were carried out using the MtrackJ plugin in ImageJ software. Dynamic parameters as migration velocity and mean square displacement (MSD) were calculated with the open-source computer program DiPer (Gorelik and Gautreau, 2014).

### Statistical analysis

All statistical tests were performed using GraphPad Prism 8 (GraphPad Software, San Diego, CA, USA).

## Notes

### Competing Interest Statement

The authors have declared no competing interest.

