## Supplemental information for "Loss of CD24 promotes radiation- and chemo-resistance by inducing stemness properties associated with a hybrid E/M state in breast cancer cells"

**SUPPLEMENTARY MATERIALS AND METHODS**

**Mammosphere formation Assay**

For mammosphere formation, cells were suspended at a concentration of 10^4^ cells/mL, in serum-free culture medium, supplemented with 1/50 B27, 20ng/mL EGF and 10ng/mL bFGF. Cells were seeded in a 6-well ultra-low adherent plate (2mL/well). Mammospheres were counted 7 days after seeding. After the culture period, mammospheres with a diameter greater than 80µm were counted. Mammosphere Forming Efficiency (MFE%) calculation:

MFE(%) = (# of mammospheres per well) / (# of cells seeded per well) x 100.

For each cell line, three independent experiments were performed.

**Irradiation**

Gamma-irradiations were performed on a GSR D1 irradiator (Gamma Medical Service). This self-shielded device irradiates with four sources of 137Cs, with a total activity around 180.28 TBq (measured in March 2014). The samples were irradiated at different single doses, namely: 0, 2, 4, 6 and 10 Gy, with a dose rate of 2.7 Gy/min, taking the radioactive decrease into account. The samples were irradiated in 25 or 75 cm^2^ flasks or 6- or 12- well - plates.

Prior to irradiation, dosimetry was performed. A cylindrical ionizing chamber 31,010 by PTW was used as the recommendation of the AAPM’S TG-61. This ionizing chamber has a cavity of 0.125 cm3 calibrated in 137Cs air kerma free in air at the PTB reference facility number 1904442. The polarity and the ion recombination were measured for this 137Cs source. Each measurement was corrected by the KTP factor to take the variation of temperature and atmospheric pressure into account.

**Primers used for quantitative real-time PCR**

All the primers used are Taqman® Assays from Applied Biosystems.

CD24: Hs02379687_s1

CD44: Hs01081473_m1

ESA (EpCAM): Hs00901885_m1

E-Cad: Hs01013965_m1

N-Cad: Hs00983056_m1

Fn1: Hs01549976_m1

Twist1: Hs00361186_m1

Twist2: Hs00382379_m1

Zeb1: Hs00232783_m1

Zeb2: Hs00207691_m1

Snai1: Hs00195591_m1

Snai2: Hs00950344_m1

Vim: Hs00185584_m1

Krt14: Hs00265033_m1

Ovol2: Hs01067398_m1

ΔNP63α: Hs00978339_m1

SOD2: Hs00167309_m1

HMOX1: Hs01110250_m1

GSR: Hs00167317_m1

TXNRD1: Hs00917067_m1

GAPDH: Hs99999905_m1

RPLPO: Hs99999902_m1

**SUPPLEMENTARY FIGURE LEGENDS**

**Supplementary Figure S1**

Analysis by qRT-PCR of the relative expression of the CD24 mRNA in E, E_CD24-, E_CD24-c and M cells. Normalization was performed as indicated in materiel and methods. Expression in E cells was normalized to 1. Each value corresponds to the mean value of 3 independent PCRs performed from 3 independent experiments. Error bars correspond to SEM.

**Supplementary Figure S2**

Representative FACS analysis of the ALDH^+^ subpopulation using the Aldefluor assay. Cells incubated with the specific inhibitor of ALDH, DEAB, were used to establish the baseline fluorescence of these cells and to define the ALDH positive population.

**Supplementary Figure S3**

Forced extinction of CD24 expression alone promotes chemo-resistance of two epithelial breast cell lines: MCF7 and T47D.

The two cell lines were transfected with the p-EBV- plasmid expressing a CD24 siRNA, in order to obtain MCF7_CD24^-^ and T47D_CD24^-^ cells. The parental and the transfected cell lines were exposed three days to 400µM 5FU and 15µM Cisplatin, and the percentage of dead cells was analyzed. Results correspond to the mean ± SD of 3 independent experiments. *P<0.05.

**Supplementary Figure S4**

Ratio of fluorescence between TMRE and Mitotracker Green analysis for E, E_CD24-, E_CD24-c and M cells.

**SUPPLEMENTARY FIGURES**

**Supplementary Figure S1**

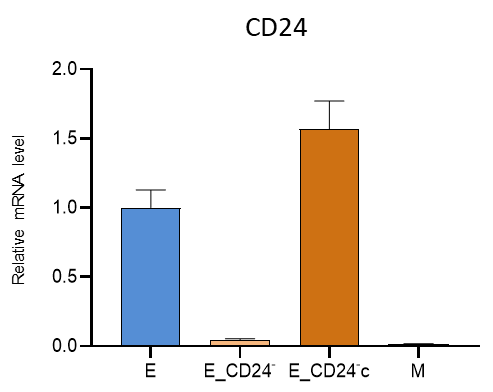

**Supplementary Figure S2**

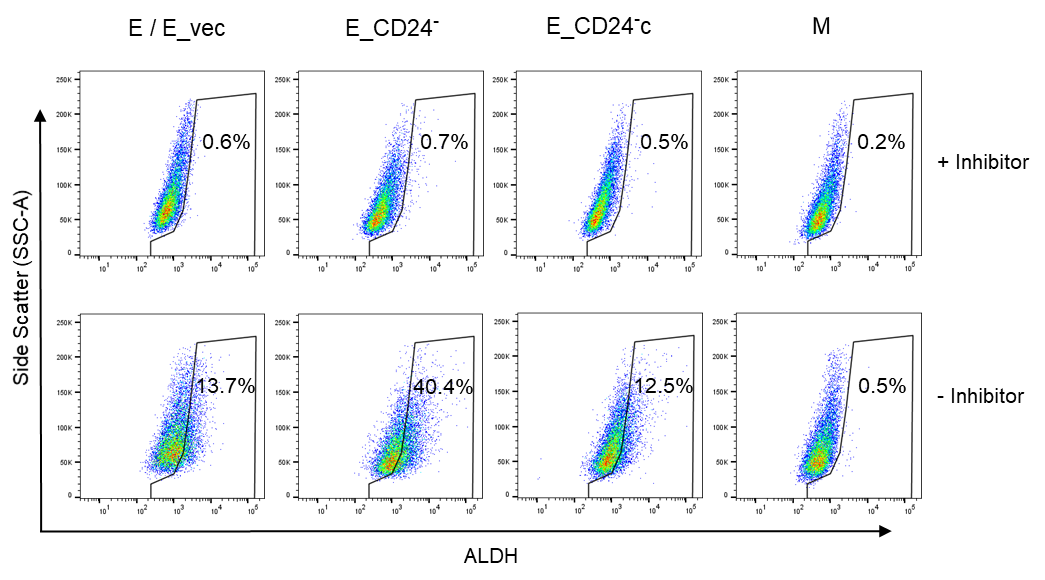

**Supplementary Figure S3**

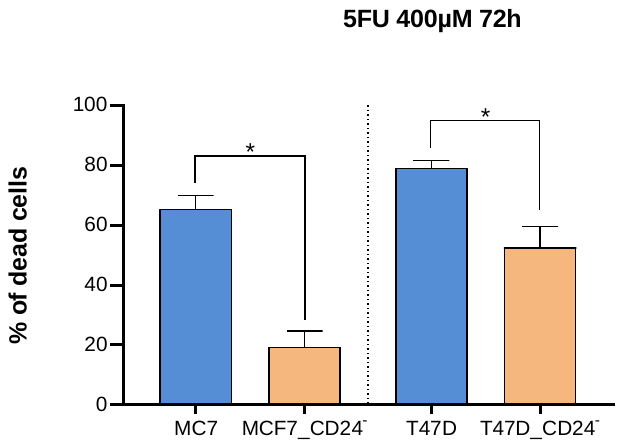

5FU (400 µM)

Day 3

% of dead cells

Cisplatine (15 µM)

Day 3

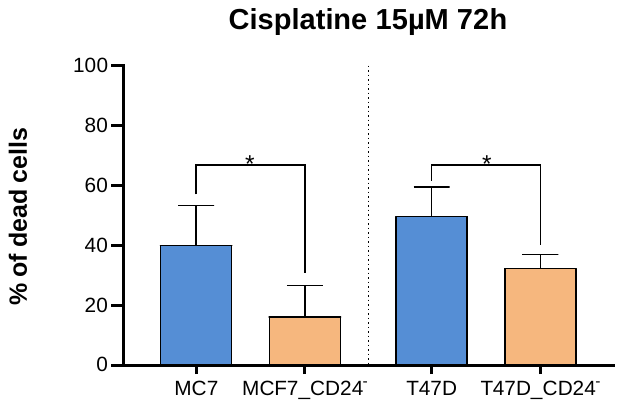

**Supplementary Figure S4**

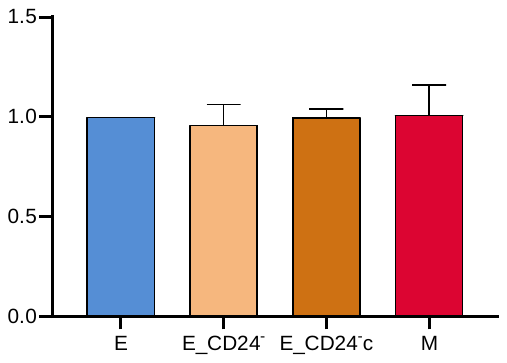

Ratio TMRE/MTG
